# A Regularized Cox Hierarchical Model for Incorporating Annotation Information in Predictive Omic Studies

**DOI:** 10.1101/2024.03.09.584239

**Authors:** Dixin Shen, Juan Pablo Lewinger, Eric Kawaguchi

## Abstract

**Background:** Associated with high-dimensional omics data there are often “meta-features” such as biological pathways and functional annotations, summary statistics from similar studies that can be informative for predicting an outcome of interest. We introduce a regularized hierarchical framework for integrating meta-features, with the goal of improving prediction and feature selection performance with time-to-event outcomes.

**Methods:** A hierarchical framework is deployed to incorporate meta-features. Regularization is applied to the omic features as well as the meta-features so that high-dimensional data can be handled at both levels. The proposed hierarchical Cox model can be efficiently fitted by a combination of iterative reweighted least squares and cyclic coordinate descent.

**Results:** In a simulation study we show that when the external meta-features are informative, the regularized hierarchical model can substantially improve prediction performance over standard regularized Cox regression. We illustrate the proposed model with applications to breast cancer and melanoma survival based on gene expression profiles, which show the improvement in prediction performance by applying meta-features, as well as the discovery of important omic feature sets with sparse regularization at meta-feature level.

**Conclusions:** The proposed hierarchical regularized regression model enables integration of external meta-feature information directly into the modeling process for time-to-event outcomes, improves prediction performance when the external meta-feature data is informative. Importantly, when the external meta-features are uninformative, the prediction performance based on the regularized hierarchical model is on par with standard regularized Cox regression, indicating robustness of the framework. In addition to developing predictive signatures, the model can also be deployed in discovery applications where the main goal is to identify important features associated with the outcome rather than developing a predictive model.

## 1 Background

Prediction based on high-dimensional omics data such as gene expression, methylation, and genotypes are important for developing better prognostic and diagnostic signatures of health outcomes. However, developing prediction models with high-dimensional omics data, where the number of features is often orders of magnitude larger than the available number of subjects is challenging. Sparse regularized regression, which includes the Lasso (1) and its variants, elastic net (2), adaptive Lasso (3), group Lasso (4) and others, is a widely used approach for developing predictive models with high-dimensional data. Sparse regularized regression controls the model complexity via sparsity inducing penalties, which have the effect of shrinking the regression coefficient estimates toward zero and setting some coefficients exactly to zero, effectively selecting features predictive of the outcome.

Associated with high-dimensional omics data, there are often prior knowledge relating to the omic features that can be informative of the outcome of interest, i.e., meta-features. For example, to predict survival based on gene expression profiles, relevant information may be the grouping of genes into biological pathways. Gene grouping information can be encoded by an indicator meta-feature matrix, each row represents one gene, each column represents one gene group, 1 indicates gene belongs to the group and 0 otherwise. This type of prior knowledge provides information on the functions of genes, available in gene annotation resources. Integrating such annotation can give omic features in these groups extra importance, via e.g., higher weights for features in these groups or shrinking weights for features outside these groups, thus improving the precision of model parameter estimation. Another important type of meta-features is summary statistics from similar studies on identical omic features. Here, the omic features could be gene expressions, and the meta-features could be regression coefficients of single nucleotide polymorphisms (SNPs) from these studies. They can fit into the same meta-feature matrix as is used to encode gene annotations, each row represents one gene expression, each column represents one SNP, the values are regression coefficients of each SNP associated with each gene. This setting is similar to transcriptome-wise associations study (TWAS), in which we investigate genetic variants’ effect on the outcome of interest through regulating expression levels. A simpler version of summary statistics is p-values, linkage disequilibrium (LD) scores, or regression coefficients from studies on the same outcome and identical omic features. The meta-feature matrix in this case only has several columns, e.g., p-values, LD scores, with each row represents respective summary statistics for a particular omic feature. This type of meta-feature is also referred to as co-data in some studies (5, 6). A third type of meta-feature is multi-omics data. In studies where data such as SNPs, gene expressions, protein levels, DNA methylations, are available, it can be insightful to integrate them altogether in one model. In this case, the meta-feature matrix is also an indicator matrix, each row represents one omic feature, each column represents one omic type (e.g., SNPs, protein levels), 1 indicates the feature belongs to a type of omic data and 0 otherwise. As more omics data and resources become available, there will be more types of data that can fit into the meta-feature framework.

Kawaguchi et. al. (2022) (7) have shown in linear regression that integrating additional prior information into regularized regression can yield improved prediction of an outcome of interest based on high-dimensional omic features. They developed a regularized hierarchical regression framework that can incorporate external meta-feature information directly into the predictive analysis with omic data. The approach is implemented in the R package xrnet. However, their method can only handle quantitative and binary outcomes. And the penalty types they are able to use are restricted to ridge. Therefore, feature selection is not available in their model. Since survival prediction is the main goal in many prognostic applications, we introduce a regularized hierarchical model, building on Kawaguchi et al. and Weaver et.al. (8), that can handle time-to-event outcomes and that can also perform meta-feature selection by the inclusion of a Lasso or elastic net regularization penalty.

There are many approaches for assessing the importance of meta-features after an analysis relating genomic features to an outcome of interest is performed. For example, gene set enrichment analysis (GSEA) (9-11) is performed after differential expression analysis to evaluate whether sets of related genes like those in the same biological pathway are over-represented.

However, there are few approaches capable of incorporating meta-features directly into the modeling process. Approaches to incorporating meta-features a priori include the application of differential penalization based on external information and two-stage regression methods, where the outcome is first regressed on the genomic features and the resulting effect estimates are in turn regressed on the external meta features. Tai and Pan (2007) (12) grouped genes based on existing biological knowledge and applied group-specific penalties to nearest shrunken centroids and penalized partial least squares. Bergerson et al. (2011) (13) incorporates external meta-feature information by weighting the LASSO penalty of each genomic feature with some function of meta-feature. Zeng et al. (2020) (14) on the other hand, incorporates external meta-feature to customize the penalty of each feature with a different function of meta-feature. These three methods are based on idea 1), which no longer assuming every genomic feature are equally important, but of different importance based on external information. However, they cannot handle large number of meta-features. Chen and Witte (2007) (15) applied the idea of hierarchical modeling in a Bayesian framework, where second stage linear regression is normal prior distribution, first stage regression is normal conditional distribution, and estimated first stage regression coefficients with posterior estimator. This method does not apply to high dimensional data since it is standard regression with no regularization. The above data integration methods improve prediction compared to modeling with genomic features only. However, none of the approaches above can handle time-to-event outcomes.

In this paper, we introduce a regularized Cox proportional hazard hierarchical model to integrate meta-features. We will see that when external meta-features are informative, regularized hierarchical modeling improves prediction performance considerably. On the other hand, we also show that when the external meta-features are not informative, it does not perform worse than the standard regularized model, that does not use any external information. This shows that the model is robust to the informativeness of the meta-features and can be safely used when the meta-feature informativeness is a priori unknown, as it is typically the case. The model can be efficiently fitted using a combination of iterative reweighted least squares and cyclic coordinate descent as proposed for Lasso Cox regression by Simon et al. (16).

## 2 Methods

### 2.1 Setup and notations

We assume a survival analysis setting with outcomes from *n* subjects, (***y***,***δ***) = (*y*_1_,…, *y*_*n*_, *δ*_1_, …,*δ*_*n*_), where ***δ*** = (*δ*_1_,…,*δ*_*n*_) is a vector of censoring status for each subject, *δ*_*i*_= 1 represents event occurs, *δ*_*i*_ = 0 represents right-censoring; ***y*** = (*y*_1_, …, *y*_*n*_) is the vector of observed time, if *δ*_*i*_ = 1, *y*_*i*_ is event time, and if *δ*_*i*_ = 0, *y*_*i*_ is censoring time. Let ***X*** denote the *n* × *p* design matrix, where *p* is the number of features, each row represents the observations on one subject, and each column represents the values of one feature across the *n* subjects. We are particularly interested in the high dimension setting, *p* ≫ *n*, where the number of features is larger than the sample size. The goal is to develop a predictive model for the outcome (***y, δ***) based on the data ***X***.

In a genomics context, the time-to-event outcome (***y, δ***) could be event free time, time to disease relapse, time to death. The design matrix ***X*** could be genotypes, gene expressions, DNA methylation. For example, in Molecular Taxonomy of Breast Cancer International Consortium (METABRIC) data (section 3.2.1), outcome (***y, δ***) is breast cancer specific survival, data matrix ***X*** represents gene expressions with dimension *number of patients* × *number of genes*.

Associated with each feature there is typically a set of meta-features annotations. If ***X*** consists of gene expression values, pathway gene sets could be meta-features indicating the set of genes involved. As for the METABRIC example, four meta-features are believed to be associated with breast cancer: genes with mitotic chromosomal instability (CIN), mesenchymal transition (MES), lymphocyte-specific immune recruitment (LYM), and FGD3-SUSD3 genes. Each meta-feature consists of a vector of indicator variables for whether a gene belongs to the functional gene group. The genomic meta-features can be collected into a matrix ***Z*** of dimensions *p* × *q*, where *q* is the number of meta-features. We propose a regularized hierarchical regression for integrating the external meta-feature information in ***Z*** for predicting time-to-event outcomes based on the features in ***X***

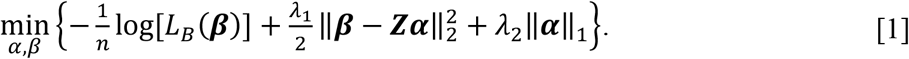

*L*_*B*_ (***β***) is the negative log of the Cox partial likelihood function, ***β*** is a length *p* vector of regression coefficients corresponding to the features in ***X***, and ***α*** is a length *q* vector of regression coefficients corresponding to the meta-features in ***Z***. The objective function [1] can be viewed as arising from a hierarchical model. In the first level of the hierarchy, the partial likelihood *L*_*B*_(***β***) term in [1] corresponds to the time-to-event outcome modeled as a function of ***X*** via a Cox proportional hazard regression model. In the second level, the *L*_2_ penalty term 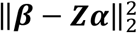 corresponds to a linear regression of the estimate of ***β*** on the meta-feature information ***Z***. It can also be thought of as an *L*_2_ regularization term that shrinks the estimate of ***β*** toward ***Zα*** rather than to the usual shrinkage toward zero. In the third level of the hierarchy, the term **‖*Z*‖**_1_ is an *L*_1_ regularization penalty on the vector of estimated effects 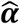. It enables the selection of important meta-features by shrinking many of its components to 0. The hyperparameters *λ*_1_, *λ*_2_ ≥ 0 control the degree of shrinkage applied to each of the penalty terms and can be tuned by cross-validation. Finally, note that when ***α*** = 0, the objective function [1] reduces to a standard *L*_2_-regularized Cox regression.

The partial likelihood function *L*_*B*_ (***β***) in [1] is the Breslow approximation (17) to the Cox partial likelihood. Letting *t*_1_ < *t*_*2*_ < … < *t*_*k*_(*k* = 1, …, *D*) be unique event times arranged on increasing order, the Cox model assumes proportional hazards:

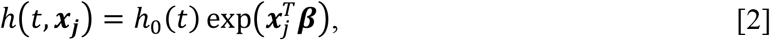

where *h*(*t*, ***x***_***j***_) is the hazard rate for subject *j* with feature values ***x***_***j***_ at time *t*; *h*_0_(*t*) is baseline hazard rate at time *t*, regardless of the feature values. The Cox partial likelihood function (18) can then be written as

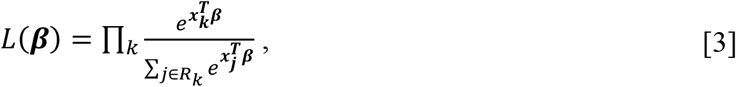

where *R*_*k*_= {*j*: *y*_*j*_≥ *t*_*k*_}, is the risk set at time *t*_*k*_, i.e., the set of all subjects who have not experienced the event and are uncensored just prior to time *t*_*k*_. The partial likelihood function allows estimation of ***β*** without explicitly modeling the baseline *h*_0_, and it depends only on the order in which events occur but not on the exact times of occurrence. However, the partial likelihood assumes that event times are unique. To handle ties, where multiple individuals experience the event at the same time, we use the Breslow approximation of the partial likelihood in [3]

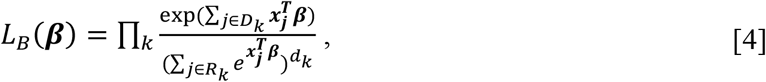

where *D*_*k*_= {*j*: *δ*_*j*_= 1, *y*_*j*_= *t*_*k*_, is the set of individuals who have event time *y*_*k*_, and *d*_*k*_ = ∑_*j*_ *I*(*δ*_*j*_ = 1, *y*_*j*_ = *t*_*k*_) is the number of events at time *y*_*k*_. Breslow’s likelihood function automatically reduces to the partial likelihood when there are no ties.

### 2.2 Computations

The objective function [1] can be minimized efficiently using iterative reweighted least squares combined with coordinate descent (16). If the current estimates of the regression coefficients are, 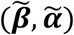 we form a quadratic approximation to the negative log-partial likelihood by Taylor series around the current estimates. The approximated objective function has the form:

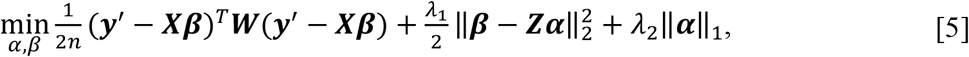

where,

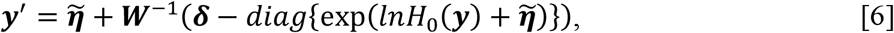

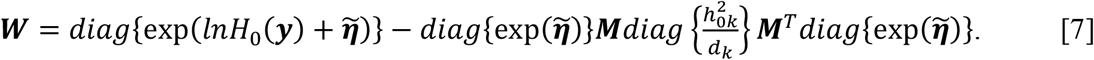

In [6] and [7], *diag* (***a***) is a diagonal matrix with vector ***a*** as diagonal elements. ***M*** is an *n* × *D* indicator matrix with (*i, k*)th element *I*(*y*_*i*_ ≥ *t* _*k*_). Also, 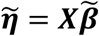 is the linear predictor; 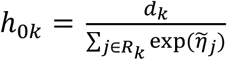 is estimated baseline hazard rate at event time *y*_*k*_ ; 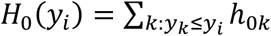 is cumulative baseline hazard at time *y*_*i*_. In the first part of quadratic approximation [5], 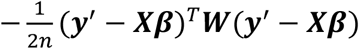 can be viewed as a weighted version of least squares with ***y***^′^ working as responses, ***W*** as weights. Weight matrix ***W*** is usually a diagonal matrix, however, in Cox proportional hazard model, ***W*** is a full symmetric matrix as shown in [7]. This leads to computational difficulty as it requires calculation of *n*^2^ entries. According to Simon et al. (16), only the diagonal entries of ***W*** are needed for computations without much loss of accuracy, thereby speeding up implementation. The diagonal elements of ***W***, *w*_*i*_ has the form:

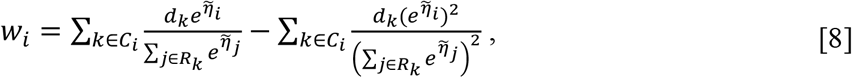

where *C*_*i*_ is the set of unique event time *t*_*k*_ such that *t*_*k*_ < *y*_*i*_ (the times for which observation *i* is still at risk). In computing weights *w*_*i*_ ’s, one bottleneck is that for each *k* in *C*_*i*_, we need to calculate 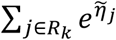. Both *C*_*i*_ and *R*_*k*_have *n* elements, so the weight computation complexity is *O*(*n*^2^). However, if *y*_*i*_ ’s are sorted in non-decreasing order, it is possible to reduce the weight computation complexity to be linear. Details are in Appendix A.1.

Now, let ***γ*** = ***β*** *−* ***Zα***, and use only diagonal elements of ***W***, the quadratic approximation [5] can be re-written as:

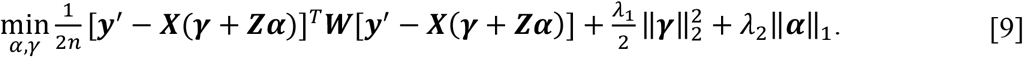

This reduced the problem to repeatedly solving the regularized, weighted least squares problem using cyclic coordinate descent (19). Details are given in Appendix A.2.

The model learning process of regularized regression is controlled by shrinkage of regression coefficients toward 0, i.e., bias, and model complexity, i.e., variance. The more shrinkage of regression coefficients toward 0, the less complex the model is, and vice versa. Further examining equation [9], the regression coefficients of omic features, ***β***, are now represented by the first level coefficients ***γ*** plus second level information ***Zα***, i.e., ***β*** = ***γ*** + ***Zα***. And ***β*** is penalized separately in ***γ*** and ***α***. Taking gene functional groups as an example for meta-feature matrix ***Z***, if a functional group is highly associated with the outcome of interest, the omic features in that group will be given extra importance by adding ***Zα***, thus coefficients of those features are less biased toward 0 (closer to unbiased maximum likelihood estimators), at the cost of added model complexity in ***α***. Now, provided the meta-features are highly informative in that unrelated functional group coefficients, a subset of ***α***, are shrunk to small values or 0 by *L*_1_ norm, the gain in bias reduction will outweigh added model complexity.

### 2.3 Two-dimensional hyperparameter tuning

The optimization approach described above is for fitting the model for one combination of the tuning parameters (*λ*_1_, *λ*_2_). More than one value combination of *λ*_1_, *λ*_2_ are usually of interest, as *λ*_1_, *λ*_2_ are tuned by cross-validation to get the best performance out of the model. For the proposed model, a two-dimensional grid of *λ*_1_, *λ*_2_ values are constructed, and pathwise coordinate optimization (20) is applied along the two-dimensional path. The pathwise algorithm utilizes current estimates as warm start, since the solutions to the convex problem [9] is continuous. This character makes the algorithm remarkably efficient and stable.

The two-dimensional hyperparameter (*λ*_1_, *λ*_2_) grid is comprised of a path of solutions corresponding to each combination of *λ*_1_, *λ*_2_. *λ*_1_ controls the amount of shrinkage to *L*_2_ term 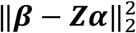, or in the transformation form in equation [9], 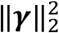. Model parameter solutions are usually initialized at 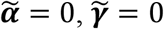. Setting the starting value of *λ*_1_, i.e., 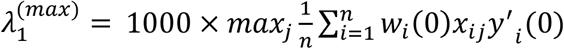, gives rise to small values of solutions, 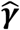. This makes convergence faster as the initial values, 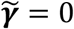, is not far away from the solutions. Applying this logic, we gradually decrease the values of *λ* and initialize the model parameters at the solutions of last *λ*, which is called warm start, until arriving at near unregularized solution. *λ*_2_, the hyperparameter controlling the amount of shrinkage to *L*_1_ term ‖α‖ _1_, is treated the same way.

The initial value of *λ*_2_, i.e., 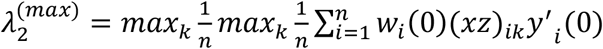, is the smallest *λ*_2_ value that makes the entire vector 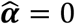. For the two-dimensional hyperparameter grid (Figure 1), we start with 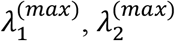, select *λ*_*min*_ = 0.01 × *λ*_*max*_, and construct a sequence of 20 *λ* values from *λ*_*max*_ to *λ*_*min*_ on log scale, forming a 20 × 20 grid with *λ*_1_, *λ*_2_ on either dimension.

**Figure 1.**
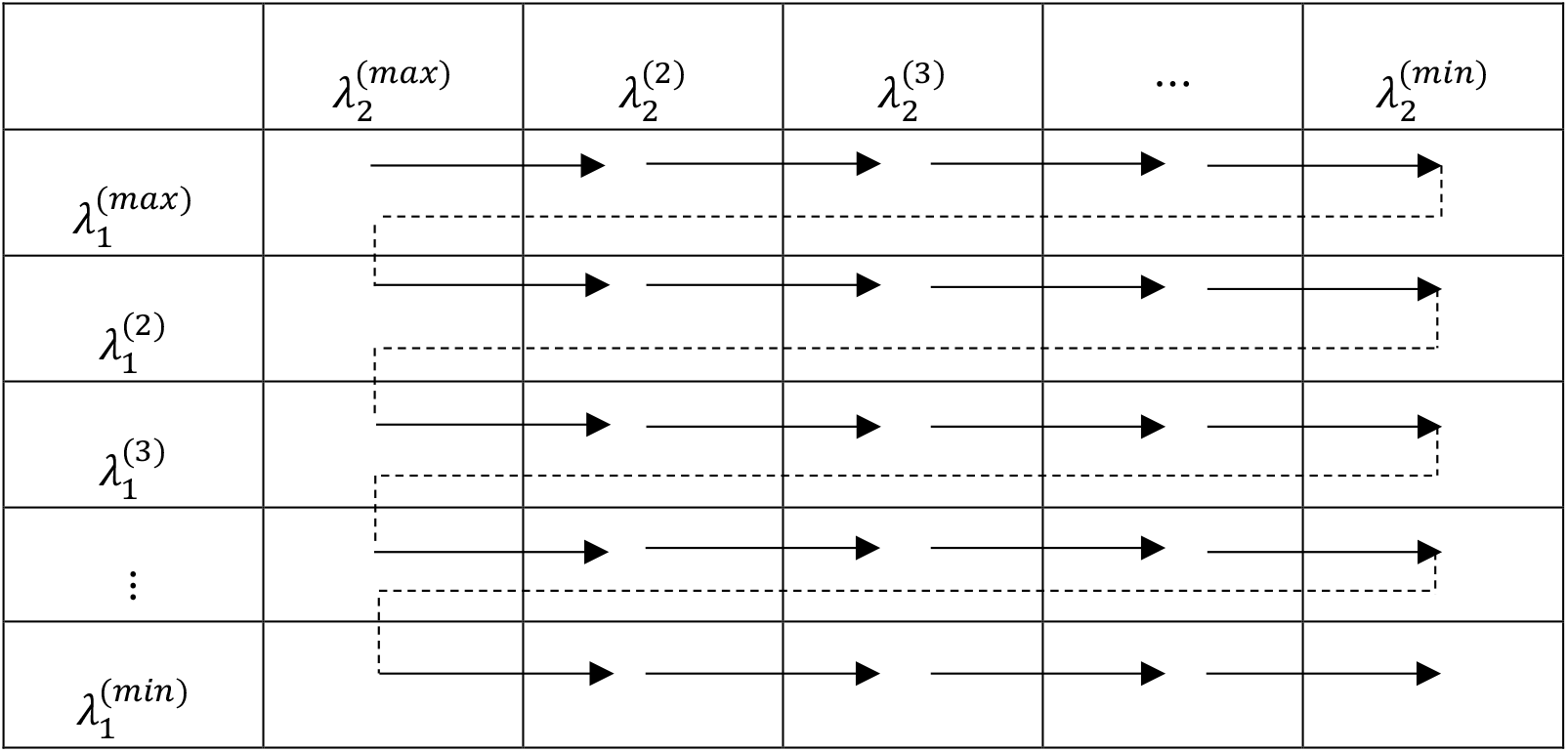
Two-dimensional hyperparameter tuning diagram

To apply warm start, one of the hyperparameters, 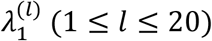, is fixed, while *λ*_2_ is decreased along the sequence until reaching 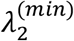. This procedure is then repeated starting at 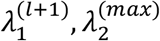,(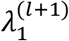 is the next value along the sequence of *λ*_1_), with warm start at solutions of 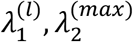. The entire 20 ***×*** 20 grid is walked through in this way and ended at 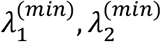.

### 2.4

#### Algorithm

The computational procedure to fit the regularized hierarchical Cox model can be summarized as:

- Initialize ***β*** and ***α*** with 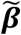 and 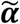.
- For each *λ*_1_, *λ*_2_, while 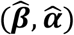 not converge:
  - Compute weights ***W*** and working response **y**^′^ with current estimate 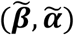, form the quadratic approximation:

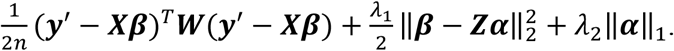
  - Find the minimizer 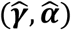 to the following optimization problem using coordinate descent. The solutions are equations [11] and [12].

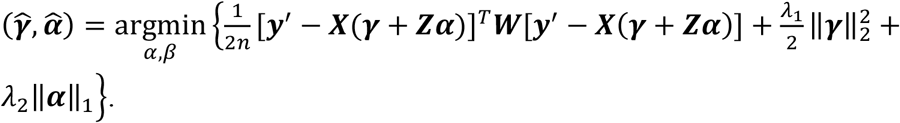
  - Set 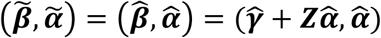.

## 3 Results

### 3.1 Simulation Study

#### 3.1.1 Simulation Design

A simulation study is performed to evaluate the predictive performance of the hierarchical Cox regression model compared to standard penalized Cox regression. The main parameters controlled include informativeness of the meta-features, sample size, number of features and number of meta-features. The *p* × *q* meta-feature matrix ***Z*** is generated with each element drawn from an independent Bernoulli variable with probability 0.1. This mimics binary indicators for whether a gene belongs to a particular biological pathway.

The first level regression coefficients are generated as ***β*** = ***Zα* + *ε***, where ***ε*** ∼ ***N***(**0**, *σ*^2^***I***). To control the predictive power of the meta-features, the signal-to-noise ratio, *SNR* = ***α***^*T*^*cov*(*Z*) ***α*** ⁄ *σ*^2^, is set, where the signal is the variance of **β** explained by ***Zα***, and σ^2^ is the noise. A higher signal-to-noise ratio implies a higher level of informativeness of the meta-features with respect to the coefficients ***β***. The data matrix ***X*** is generated by sampling from a multivariate normal distribution, *N*(0, **Σ**), where the covariance matrix **Σ** has an autoregressive correlation structure **Σ**_*ij*_ = *ρ*^|*i−j*|^ for *i, j* = 1, …, *p*.

The cumulative distribution function of the Cox proportional hazard model is given 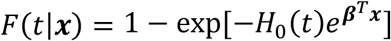, where *H*_0_(*t*) is baseline cumulative hazard function. Using the inverse probability integral transform (21), survival times *p* are generated as:

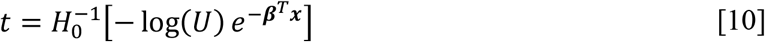

where *U*∼*uniform*[0,1]. A Weibull distribution is used for the baseline hazards, which has cumulative hazard function 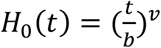. The baseline Weibull parameters are set to *b* = 5, *v* = 8, which result in survival times in the range 0 to 20. The censoring time, *c*, is simulated based on an exponential distribution with density *f*(*t*) = exp(−*λt*), with *λ* = 0.06. Then, the time-to-event outcome (*y*_*i*_, *δ*_*i*_) is generated as (min(*t*_*i*_, *c*_*i*_), *I*(*t*_*i*_ < *c*_*i*_)). The value of exponential distribution parameter *λ* is chosen to result in a ratio of subjects experiencing the event vs. subjects experiencing censoring of about 2 to 1.

To control the predictivity of the features ***X*** for the outcome ***y***, Harrell’s concordance index (C-index) (22) is used as the metric to evaluate prediction performance. It is defined as the probability that a randomly selected patient who experienced an event has a higher risk score, ***β***^*T*^***x***, than a patient who has not experienced an event at a given time. The C-index is an analog of the area under the ROC for time-to-event data. The higher the C-index, the better the model can discriminate between subjects who experience the outcome of interest and subjects who do not or have not yet. A random noise is added to the survival times *t* to control the C-index, where the noise is distributed as a normal with mean zero and a variance value set to yield a C-index of 0.8 across all simulation scenarios. This is the population/theoretical C-index of the generated survival data, achievable if ***β*** were known or if one had an infinite sample size. When ***β*** is estimated from a finite training set, the achieved model C-index will be lower.

The base case scenario is simulated with sample *N* = 100, number of features *p* = 200, and number of meta-features *q* = 50. This is a high dimensional setting, *p* ≫ *N*, typical of genomic studies. The first 20% of the coordinates of the meta-feature level coefficients ***α*** are set to be 0.2, and the rest are set to be 0. In the base scenario, the meta-features are highly informative, with a signal noise ratio set to 2. The covariance matrix **Σ** of ***X*** has autoregressive-1 structure, with parameter *ρ* = 0.*5*. In the following simulation situations, one of the parameters is varied and the others are fixed in each scenario. Simulations are performed 100 times for all scenarios. The models are trained on a training set of size *N*(100 in the base scenario and varied in other experiments), with the hyper-parameters *λ*_1_, *λ*_2_ tuned on an independent validation set of the same size as training set. The final predictive performance was evaluated on a large test set of size 10,000.

We run a series of experiment varying one key parameter at a time from the base case scenario as follows:

Experiment 1: varying the signal-to-noise ratio of the meta-features from completely uninformative, (SNR=0), to slightly informative (SNR=0.1), to moderately informative, (SNR= 1), to highly informative (SNR=2).

Experiment 2: Varying the sample size from low to high, *N* = 100, 200, 500.

Experiment 3: Varying the number of features from low to high: *p* = 200, 1000, 10000.

Experiment 4: Varying the number of meta-features from low to high: *q* = 20, 50, 100.

#### 3.1.2 Simulation results

The results of the experiments are shown in Figure 2. In each panel, the horizontal dashed line representing the population/theoretical C-index, 0.8, i.e., the maximum achievable with infinite training data, is provided as a reference for each parameter setting. We compared the performance of the hierarchical ridge-lasso Cox model (ridge penalty on first level omic features, Lasso penalty on second level meta-features) incorporating meta-features to that of a standard ridge Cox model, and the performance of the hierarchical lasso-lasso Cox model (Lasso penalty on first level omic features, Lasso penalty on second level meta-features) to that of a standard Lasso Cox model.

**Figure 2.**
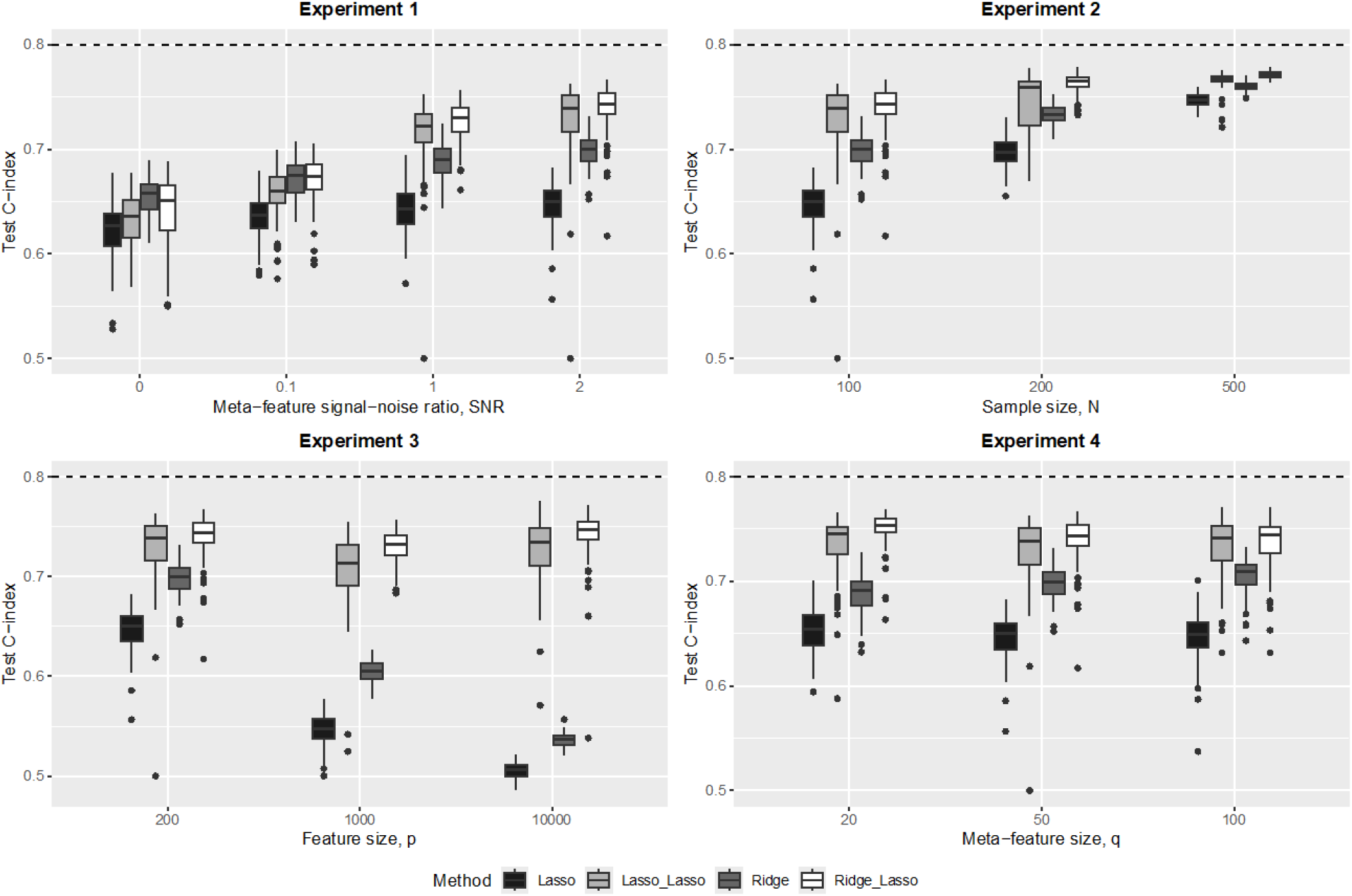
Simulation results: prediction performance. Figure 2: Prediction performance comparison between the proposed regularized Cox hierarchical model and the standard regularized Cox model, i.e., lasso vs. lasso-lasso, ridge vs. ridge-lasso. The horizontal dashed line is the theoretical C-index at 0.8 for generated datasets. The boxplot represents medium and interquartile C-index over 100 simulations. Experiment 1: N = 100, p = 200, q = 50, SNR = 0, 0.1, 1, 2. Experiment 2: N = 100, 200, 500, p = 200, q = 50, SNR = 2. Experiment 3: N = 100, p = 200, 1000, 10000, q = 50, SNR = 2. Experiment 4: N = 100, p = 200, q = 20, 50, 100, SNR = 2.

With informative meta-features (SNR > 0 in experiments 1-4) the hierarchical ridge-lasso model consistently outperforms the standard ridge model, with the performance gain over the standard ridge model increasing with the informativeness of the meta-features (experiment 1).

Importantly, when the meta-features are completely uninformative, the hierarchical ridge-lasso model performs only slightly worse than the standard ridge model (experiment 1, SNR=0). This shows robustness of the hierarchical ridge-lasso model to uninformative meta-features.

Experiment 2 shows that the gains in performance of the hierarchical ridge-lasso model over the standard ridge model can be quite large, particularly when the sample size is small. As the sample size *N* increases, the performance of both models increases and the difference between the two is reduced.

As the dimensionality *p* of the features increases (experiment 3), the performance of the standard ridge model deteriorates dramatically, while the performance of the hierarchical ridge-lasso model only decreases slowly as the information in the meta-features helps stabilize its performance.

In experiment 4, the performance of the standard ridge model does not change, as it does not utilize meta-feature information. However, for the hierarchical ridge-lasso model, the performance decreases as the number of noise meta-features increases (the number of informative meta-feature is set at 20% of the coordinates of ***α*** and the additional meta-features are noise meta-features).

The comparison between lasso-lasso model and standard Lasso model shares similar trends as those between ridge-lasso model and standard ridge model, except that they have lower prediction performances than their respective counterparts. This is not surprising that the Lasso excels in producing interpretable models, while the ridge does well in prediction.

We also examined the ability of the model to select informative meta-features by second-level Lasso penalty. In particular, we looked at the overall accuracy (percentage of alphas correctly shrunk to 0 or not shrunk to 0), true positive rate (percentage of non-zero alphas that are not shrunk to 0), and false positive rate (percentage of zero alphas that are not shrunk to 0) in experiment 1, where the second level meta-features informativeness varies (Figure 3). We see that as the informativeness of the meta-features increases, the overall accuracy increases, with the true positive selection rate of meta-features improves dramatically at the cost of a slight increase in the false positive rate.

**Figure 3.**
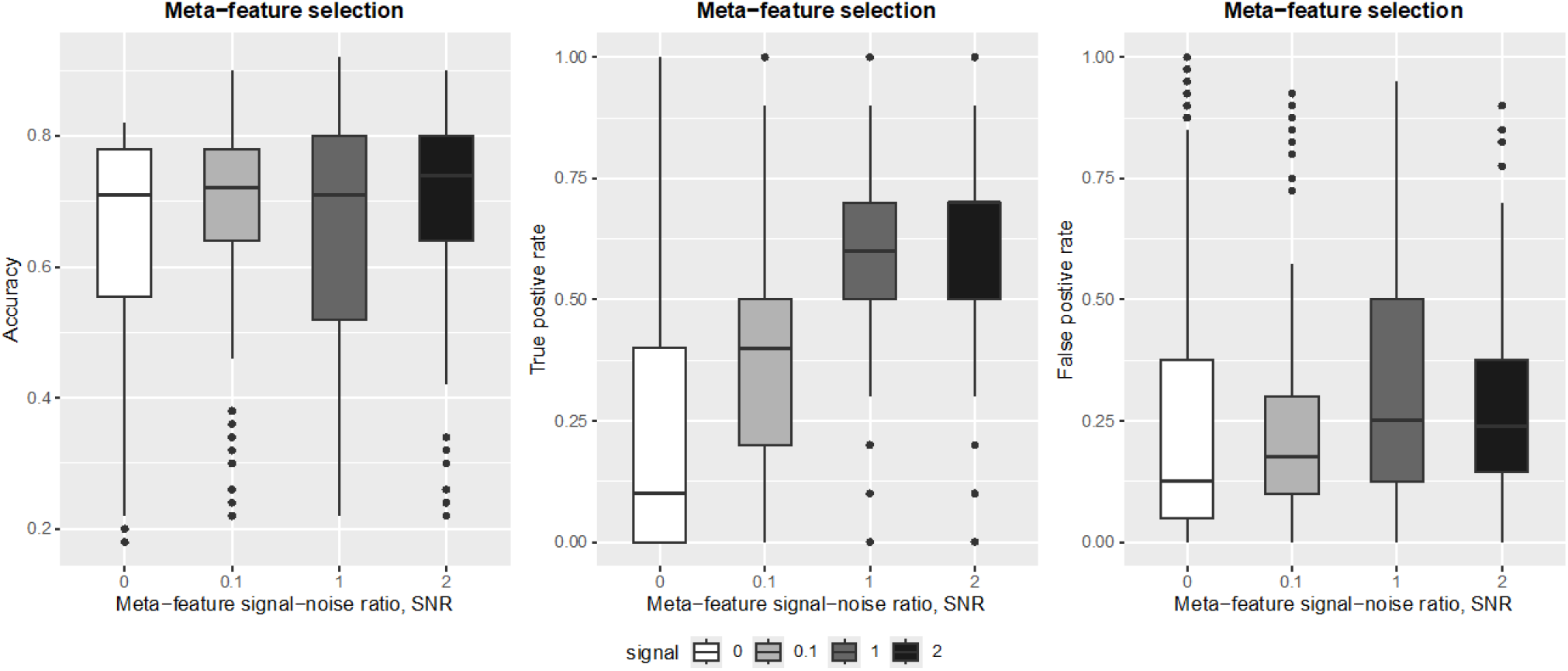
Simulation results: meta-feature selection. Figure 3: Meta feature selection performance in increasing informativeness of meta features, i.e., SNR = 0, 0.1, 1, 2, while N = 100, p = 200, q = 50. The boxplot represents median and interquartile of meta-feature selection overall accuracy, true positive rate and false positive rate over 100 simulations.

### 3.2 Applications

#### 3.2.1 Gene Expression Signatures for Breast Cancer Survival

To illustrate the performance of our approach, we applied the hierarchical survival model to the Molecular Taxonomy of Breast Cancer International Consortium (METABRIC) study (23). The METABRIC microarray dataset is available at European Genome-Phenome Archive with the accession of EGAS00000000083. It includes cDNA microarray profiling of around 2000 breast cancer specimens processed on the Illumina HT-12 v3 platform (Illumina_Human_WG-v3). The dataset was divided into a training set of 997 samples, and a test set of 995 samples (24). The goal is to build a prognostic model for breast cancer survival, based on gene expressions and clinical features. The data ***X*** consists of 29,477 gene expression probes and two clinical features, age at diagnosis and the number of positive lymph nodes. The meta-feature data ***Z*** consists of four “attractor metagenes”, gene co-expression signatures that are shared across many cancer types and are associated with specific cancer phenotypes. The shared features in cancer include, e.g., the ability of cancer cells to divide uncontrollably, to invade surrounding tissues, and, with the effort of the organism to fight cancer with a particular immune response (25). Three of the universal “attractor metagenes”, mitotic chromosomal instability (CIN), mesenchymal transition (MES), lymphocyte-specific immune recruitment (LYM), were found associated with prognosis of breast cancer. In addition, a metagene whose expression is associated with good prognosis and that contains the expression values of two genes—FGD3 and SUSD3. The CIN, MES, and LYM metagenes each consist of 100 genes, but for our analysis, we only considered the 50 top-ranked genes. The data matrix ***Z*** is an indicator matrix of whether a specific expression probe corresponds to a gene in a metagene.

Model building was based on the samples with ER positive and HER2 negative, as treatments are homogeneous in this group, and they are associated with good prognosis (26). There were 740 samples in the training set and 658 samples in the test set in the ER+ and HER2-subset after removing samples with missing values. We used repeated 5-fold cross validation to tune the hyper-parameters *λ*_1_, *λ*_2_ in the training set, with 50 repetitions. The test set was used to evaluate model performance. The same training/test scheme was used to fit a standard ridge regression without attractor metagene information as comparison.

With only gene expression features in the model and no clinical features, the mean test C-index for the ridge-lasso hierarchical model with metagene information was 0.658 (95% CI: 0.639, 0.677) which compares favorably with the mean test C-index of 0.639 (95% CI: 0.628, 0.650) for the standard Cox ridge counterpart. When adding the clinical features: age at diagnosis and number of positive lymph nodes, the mean test C-index increased to 0.734 (95% CI: 0.716, 0.752), and 0.728 (95% CI: 0.726, 0.730) for the Cox hierarchical model, and the standard Cox ridge model, respectively (Table 1). The metagenes CIN and FGD3-SUSD3 were identified by the hierarchical model as being important (had higher absolute values of coefficients, ***α***).

**Table 1.** Test C-index between standard ridge and ridge-lasso.

|  |  | Standard ridge,<br>mean (95% CI) | Ridge-Lasso,<br>mean (95% CI) |
| --- | --- | --- | --- |
| **Test C-index** | Gene expression features only | 0.639 (0.628, 0.650) | 0.658 (0.639, 0.677) |
|  | Gene expression + clinical | 0.728 (0.726, 0.730) | 0.734 (0.716, 0.752) |

Metagene CIN, which is a breast cancer inducing metagene, had a positive coefficient, indicating genes in CIN had an overall increased risk over other genes, while the FGD3-SUSD3 metagene had a negative coefficient estimate, indicating FGD3 and SUSD3 had a reduced risk (Table 2). The identified metagenes were also found by previous analysis (24).

**Table 2.**
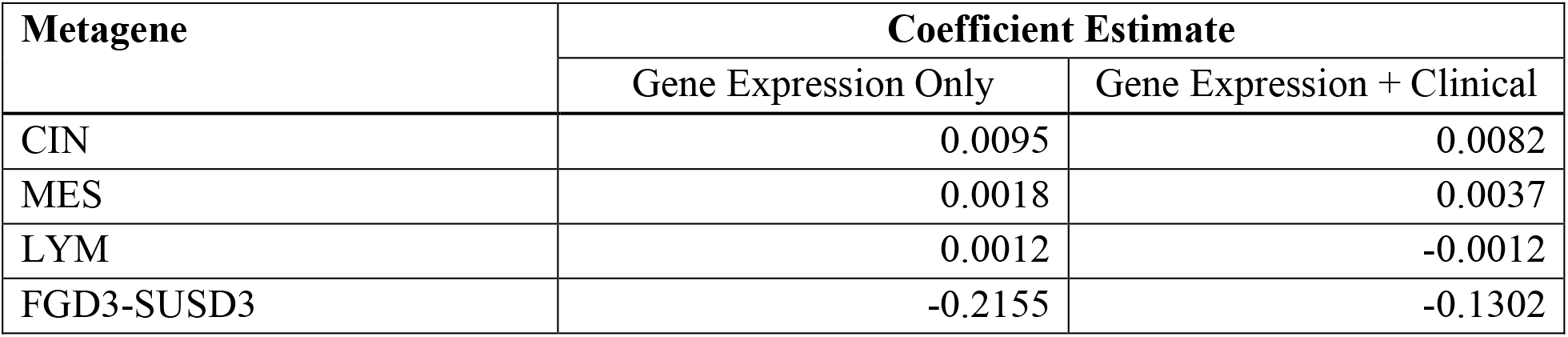
Coefficient estimates for “attractor metagenes”.

| **Metagene** | **Coefficient Estimate** |  |
| --- | --- | --- |
|  | Gene Expression Only | Gene Expression + Clinical |
| CIN | 0.0095 | 0.0082 |
| MES | 0.0018 | 0.0037 |
| LYM | 0.0012 | -0.0012 |
| FGD3-SUSD3 | -0.2155 | -0.1302 |

#### 3.2.2 Anti-PD1 Immunotherapy Predictive Biomarker for Melanoma Survival

We also applied the model to a melanoma data set to predict overall survival after treating patients with a PD-1 immune checkpoint blockade. The programmed death 1 pathway (PD-1) is an immune-regulatory mechanism used by cancer to hide from the immune system. Antagonistic antibodies to PD-1 pathway and its ligands, programmed death ligand 1 (PD-L1), has demonstrated high clinical benefit rates and tolerability. Immune checkpoint blockades such as Nivolumab, pembrolizumab are anti-PD-1 antibodies showing improved overall survival for the treatment of advanced melanoma. However, less than 40% of the patients respond to the treatments (27). Therefore, predicting treatment outcomes, identifying predictive signals are of great interest to appropriately select patients most likely to benefit from anti-PD-1 treatments.

We explored transcriptomes and clinical data using our model to illustrate prediction performance and predictive signal selection.

The dataset combined 3 clinical studies in which RNA-sequencing were applied to patients treated with anti-PD1 antibodies, Gide et al., 2019 (28), Riaz et al., 2017 (29), Hugo et al., 2016 (30). The gene expression values are normalized toward all sample average in each study as the control, so that they are comparable to one another across features within a sample and comparable to one another across samples. There are 16010 genes in common across 3 studies and 117 subjects combined. We build predictive models in terms of overall survival, based on gene expression profile. Since the subjects are all treated with anti-PD1 antibodies, the transcriptomic features selected by the model are predictive signals for treatment efficacy or resistance. We selected meta-features from molecular signature database, hallmark gene sets. The hallmark gene sets involve biological pathways such as signaling, immune, proliferation. They have been applied to analyses of cancer phenotypes of Medulloblastoma, Glioblastoma, and protein levels (31). 13 gene sets are enriched to have false positive rates less than 0.25. An indicator matrix ***Z*** is formed to illustrate whether each of the 16010 genes belong to one of the 13 hallmark gene sets.

We performed 5-fold cross validation to tune the hyper parameters and report the validation prediction performance. We see an improvement in prediction with the hallmark gene set information with a C-index of 0.663 for ridge-lasso compared to 0.637 for standard ridge. At the gene set level, 3 gene sets have absolute effect size larger than 0.01 (Table 3). Specifically, genes in response to interferon gamma, genes that are involved in KRAS regulation were identified. A subset of the genes in the identified gene sets by our model were in concordance with the previously published anti-PD1 gene signatures (29, 30).

**Table 3:** Coefficient estimates (non-zero, selected) for gene sets.

| Gene Set | Coefficient Estimate |
| --- | --- |
| IFNG interferon gamma response | -0.0100* |
| Interferon alpha response | -0.0013 |
| IL-2_STAT5 signaling | 0.0072 |
| Bile acid metabolism | -0.0011 |
| KRAS signaling down regulated | -0.0135* |
| KRAS signaling up regulated | 0.0100* |
| Apoptosis | 0.0004 |
| Xenobiotic metabolism | 0.0013 |

\* Gene sets with absolute value of coefficients larger than 0.01

## 4 Discussion

In this paper we extended the regularized hierarchical regression model of Kawaguchi et al. to time-to-event data and to accommodate a Lasso or elastic-net penalty in the second level of the model. The hierarchical regularized regression model enables integration of external meta-feature information directly into the modeling process. We showed that prediction performance improves when the external meta-feature data is informative. And the improvements are largest for smaller sample sizes, when prediction is hardest and performance improvement is most needed. Key to obtaining performance gains though is prior knowledge of external information that is potentially informative for the outcome. For example, clinicians, epidemiologists, or other substantive experts may provide insights into what type of annotations are likely to be informative. However, the model is robust to incorporating a set of meta-features that is completely irrelevant to the outcome of interest. In this scenario, a very small price in prediction performance is paid relative to a standard ridge model (i.e., without external information). This should encourage the user to integrate meta-features even if uncertain about their informativeness.

An underlying assumption of the proposed regularized hierarchical model is that the effects in a group determined by meta-features (e.g., genes in a pathway) are mostly in the same direction. A limitation of the method is that if the effects have opposite signs and ‘cancel each other out’ there would be little or no improvement in prediction, even if the pathway information is informative.

In addition to developing predictive signatures, the model can also be deployed in discovery applications where the main goal is to identify important features associated with the outcome rather than developing a predictive model. However, there is no standard way to perform formal inference, i.e., standard errors, p-values, confidence intervals, with high-dimensional regression models. Several approaches exist (32, 33) and this is an active area of research. Adding formal statistical inference would be an important future work to expand the range of use of the proposed model.

The regularized hierarchical Cox model is implemented in xrnet package and available to install via GitHub (34). The average computation times (mean computation time over 100 repetitions) of the situations in simulation experiment 3, N = 100, p = 200, 1000, 10000, q = 50, i.e., running 20 × 20 hyperparameter grid, are 0.9, 3.6, 57.4 seconds, respectively. The implementation was conducted on 8-core M2 CPU, with the operation system MAC OS 14.5. The implementation is efficient and can be used to perform analyses with large number of features, meta-features, and subjects, as is the case in METABRIC and anti PD-1 data applications in section 3.2. While the models we focused on in the simulation and data applications are mainly ‘ridge-lasso’, i.e., with an *L*_2_ norm penalty applied to ***β*** − ***Zα***, and an *L*_1_ norm applied to the meta-feature coefficients ***α***, the implementation offers the flexibility of using the Lasso, elastic net, and ridge penalties to penalize the meta-features depending on the application. As a result, different combinations of penalty types can be tuned to achieve optimal prediction performance, as well as tailored penalty types to the need of prediction or feature selection. For example, if selection at the meta-feature level is desired and the meta-features are highly correlated, the elastic net penalty is a better option for ***α*** regularization. Because if there is a group of variables that are highly correlated, the lasso tends to select one of them randomly, while the elastic net enjoys grouping effect which selects all the variables in a group with estimated coefficients close in magnitude (2). The approach does not perform feature selection on first level information as it uses a ridge penalty. In a high dimensional setting, standard regularized regression like the Lasso and elastic net often selects relatively large number of features. It can then be valuable to identify groups of genes defined by meta-features that may jointly have significant predictive power for the outcome of interest. Another potential improvement of the model is to extend the range of penalty types to non-convex penalties, such as SCAD (35), MCP (36). These penalties yield less biased effect size estimates than that of the Lasso and elastic net.

## 5 Conclusions

The proposed hierarchical regularized regression model enables integration of external meta-feature information directly into the modeling process for time-to-event outcomes. Its prediction performance improves when the external meta-feature data is informative. Importantly, when the external meta-features are uninformative, the prediction performance based on the regularized hierarchical model is on par with standard regularized Cox regression, which should encourage the user to integrate meta-features even if uncertain about their informativeness. In addition to developing predictive signatures, the model can also be deployed in discovery applications where the main goal is to identify important features associated with the outcome rather than developing a predictive model. The developed R package written with C++, xrnet, is computationally efficient, accommodates large and sparse matrices, offers the flexibility of using the Lasso, elastic net, and ridge penalties to both omic features and meta-features.

## A Appendix

### A.1 Computation of diagonal elements of weight matrix

Diagonal elements of weight matrix ***W***, the Hessian of log Cox’s partial likelihood function, *w*_*i*_, has the form:

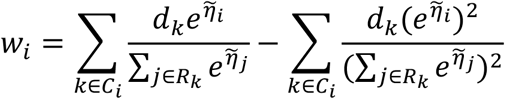

The two sums in sets *R*_*k*_ and *C*_*i*_ both have *n* elements, hence it is a *O*(*n*^2^) computation. However, if we notice the difference between the two sets *R*_*k*_ and *R*_*k*+1_is the observations that are in *R*_*k*_but not in *R*_*k*+1_, i.e., {*j*: *t*_*k*_ ≤ *y*_*j*_ < *t*_*k*+1_ }, provided that the observed times ***y*** are sorted in non-decreasing order, then 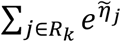 can be calculated as cumulative sums:

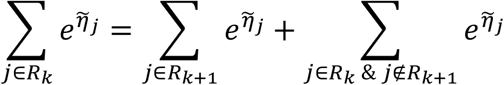

The same cumulative sum idea can be applied to the set *C*_*i*_ :

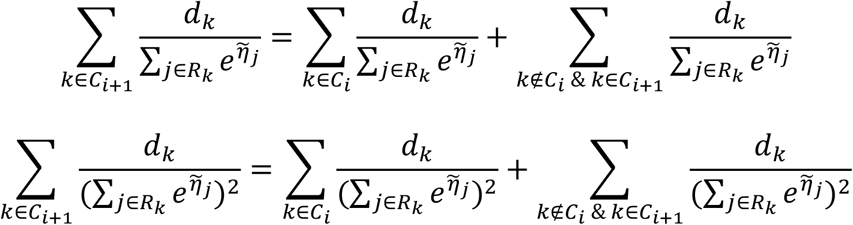

The equations above only calculate the sum once, then add at each sample index, which brings down the computation cost to linear time, *O*(*n*). Considering sorting observed times as a data pre-processing procedure, the overall computation time for the weights is *O*(*n* log *n*).

### A.2 Solve regularized weighted least squares with cyclic coordinate descent

To solve regularized weighted least squares, equation [9], We first compute the gradient at current estimates 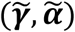. Let *γ*_*j*_ be the *jth* coordinate of ***γ***, 1 ≤ *j* ≤ *p*; *α*_*k*_ be the *kth* coordinate of ***α***, 1 ≤ *k* ≤ *q*. The gradient of equation [9] with respect to *γ*_*j*_ is

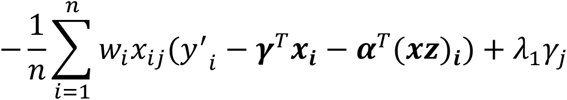

Setting the gradient with respect to *γ*_*j*_ to 0, the coordinate-wise update for *γ*_*j*_ has the form:

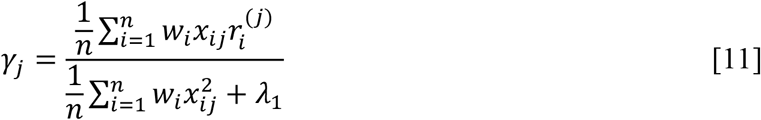

where 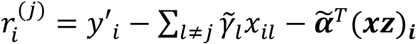, is the partial residual excluding the contribution of *x*_*ij*_. As for *α*_*k*_, if 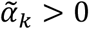, then the gradient of (9) with respect to *α*_*k*_ is

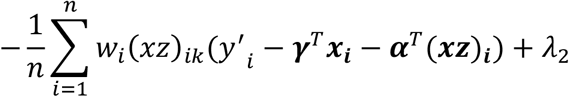

A similar expression exists if 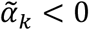, and 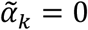 is treated separately. Setting the gradient with respect to *α*_*k*_ to 0, the coordinate-wise update for *α*_*k*_ has the form:

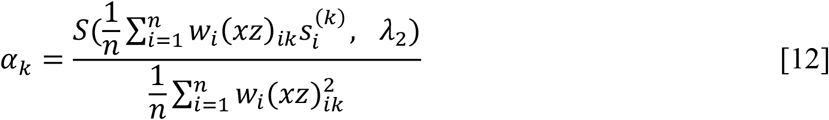

where 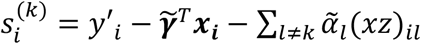, is the partial residual excluding the contribution of (*xz*)_*ik*_. *S*(*z, λ*) is the soft-thresholding operator with value:

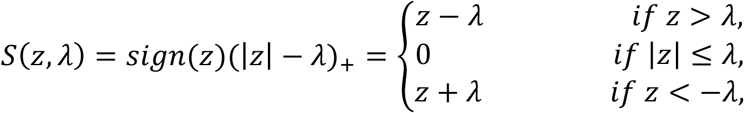

## Declarations

## I. Ethics approval and consent to participate

Not Applicable

## II. Consent for publication

Not Applicable

## III. Availability of data and materials

Source codes for simulations, section 3.1, are available at https://github.com/dixinshen/Simulation-and-Application-Data-of-Regularized-Cox-Hierarchical-Model (37).

METABRIC (23) associated genotype and expression data are available at the European Genome-Phenome Archive under accession number EGAS00000000083.

Anti-PD1 immunotherapy predictive biomarker for melanoma survival application:

i. Data associated with Gide et al., 2019 (28) are available at the European Nucleotide Archive under accession number PRJEB23709.
ii. Data associated with Riaz et al., 2017 (29) are available at the NCBI Gene Expression Omnibus under series number GSE91061.
iii. Data associated with Hugo et al., 2016 (30) are available at the NCBI Gene Expression Omnibus under series number GSE78220.

## IV. Competing interests

The authors declare that they have no competing interests.

## V. Funding

This work was supported by the National Cancer Institute at the National Institutes of Health [P01 CA196569, R01 CA140561].

## VI. Authors’ contributions

D.S. developed the methodology, programmed the implementation, performed simulations and applications, drafted and reviewed the manuscript. J.P.L. co-developed the methodology, acquired METABRIC data, reviewed the manuscript. E.K. programmed the implementation.

## VII. Acknowledgements

Not Applicable

## Notes

### Competing Interest Statement

The authors have declared no competing interest.

### Summary of Updates

Add an author, Eric Kawaguchi, Update Author's contributions.

